# New role for cardiomyocyte *Bmal1* in the regulation of sex-specific heart transcriptomes

**DOI:** 10.1101/2024.04.18.590181

**Authors:** Xiping Zhang, Spencer B. Procopio, Haocheng Ding, Maya G. Semel, Elizabeth A. Schroder, Tanya S. Seward, Ping Du, Kevin Wu, Sidney R. Johnson, Abhilash Prabhat, David J. Schneider, Isabel G Stumpf, Ezekiel R Rozmus, Zhiguang Huo, Brian P. Delisle, Karyn A. Esser

**Author notes:** These authors contributed equally to this paper (Xiping Zhang, Spencer B. Procopio).

## Abstract

It has been well established that cardiovascular diseases exhibit significant differences between sexes in both preclinical models and humans. In addition, there is growing recognition that disrupted circadian rhythms can contribute to the onset and progression of cardiovascular diseases. However little is known about sex differences between the cardiac circadian clock and circadian transcriptomes in mice. Here, we show that the the core clock genes are expressed in common in both sexes but the circadian transcriptome of the mouse heart is very sex-specific. Hearts from female mice expressed significantly more rhythmically expressed genes (REGs) than male hearts and the temporal pattern of REGs was distinctly different between sexes. We next used a cardiomyocyte-specific knock out of the core clock gene, *Bmal1*, to investigate its role in sex-specific gene expression in the heart. All sex differences in the circadian transcriptomes were significantly diminished with cardiomyocyte-specific loss of *Bmal1*. Surprisingly, loss of cardiomyocyte *Bmal1* also resulted in a roughly 8-fold reduction in the number of all the differentially expressed genes between male and female hearts. We conclude that cardiomyocyte-specific *Bmal1*, and potentially the core clock mechanism, is vital in conferring sex-specific gene expression in the adult mouse heart.

## Introduction

Sex differences are ubiquitous in mammals at the organism, tissue, and gene expression levels. The heart exhibits sex differences in many aspects, including structure, function, and differential responses to disease and drug treatment.^1,2^ For example, in humans, males tend to have larger hearts, greater cardiac output, and a lower heart rate compared to females.^3^ Sex-specific gene expression contributes to these morphological and functional differences and has been observed in humans and mice.^4–7^ Multiple factors, including sex chromosomes and sex hormones, drive these transcriptional differences. While most ubiquitously expressed transcription factors are not differentially expressed between males and females, they exhibit transcriptional regulation in a sex-specific manner. As a result, genes that are differentially expressed between males and females have been associated with sex-specific function and disease in the heart.^6,8^

Some of the most ubiquitosly expressed transcription factors are those responsible for regulating circadian gene expression. These intrinsic circadian oscillations in gene expression produce 24-hour rhythms in physiology, metabolism, and behavior, allowing organisms to anticipate and adapt to daily environmental changes. At the molecular level, the core clock mechanism is described as a transcriptional and translational feedback mechanism in which the transcriptional activators BMAL1 and CLOCK heterodimerize and bind to Ebox-containing genes and induce the expression of the negative limb genes, including *Per1/Per2, Cry1/Cry2*, and *Rev-erb*s. Beyond timekeeping, the clock mechanism also contributes to a daily program of gene expression, termed clock output. While the activator-repressor core clock exists in all cells, the clock output genes differ between tissues and function to coordinate time-of-day tissue-specific physiology.^1,2,9,10^ Several studies have demonstrated the importance of the clock mechanism in the heart as alterations result in tissue physiological and functional changes.^11–14^

Sex differences in the circadian transcriptome have recently been reported in mouse liver, in which more rhythmically expressed genes (REGs) were identified in females than in males.^15,16^ A similar result was also reported in a human study, in which the authors used Genotype-Tissue Expression (GTEx) with an algorithm to assign circadian phase. This work found that females displayed more REGs than males in multiple tissues. Of those REGs identified in the heart, only 50% were shared between sexes, while others were unique to one sex or showed sex-specific rhythmic patterns.^17^ However, little work has been done to parse out these differences in rhythmic gene expression and to determine the contribution of the core clock mechanism to this phenomenon.

In this study, we investigated the role of cardiomyocyte-specific *Bmal1* on the cardiac circadian transcriptome of adult male and female mice using RNAseq. Consistent with other studies, we observed that female hearts express a significantly greater number of REGs compared to male hearts, with a modest overlap in genes and functional enrichment. Analysis of the temporal distribution of the REGs, using peak expression time, illustrated that the distribution of REGs throughout the day was completely different between sexes. Most REGs in the male heart peak at the transition from rest to active or active to rest phases. In contrast, most REGs in the female heart peak in the middle of the rest or middle of the active periods. Surprisingly, cardiomyocyte-specific loss of *Bmal1* resulted in a significant reduction of all sex differences in the circadian transcriptome. In addition, analysis of all differentially expressed genes (DEGs), including non-rhythmic genes, between males and females showed that differential expression between sexes was largely diminished with loss of cardiomyocyte *Bmal1*. We conclude that cardiomyocyte-specific *Bmal1*, and likely the core clock mechanism, is vital in conferring a sex-specific gene expression program in the adult mouse heart.

## Methods

All animal procedures were performed by the Association for Assessment and Accreditation of Laboratory Animal Care guidelines and approved by the University of Kentucky Institutional Animal Care and Use Committee and the University of Florida Institutional Animal Care and Use Committee.

The inducible cardiomyocyte-specific *Bmal1* knockout mice (iCS*Bmal1* KO) were bred by crossing the floxed *Bmal1* mouse with the cardiomyocyte-specific Myh6-MerCreMer recombinase transgenic mouse as described previously.^14^ All mice were born from the same breeders and housed in the same facility for the total duration of the experiment. Cre recombination was induced at 3-4 months of age by intraperitoneal injection of tamoxifen (35mg/kg/day) for three consecutive days to induce the deletion of *Bmal1* in adult cardiomyocytes. The control mice were generated by injecting Cre^+/-^ *Bmal1*f/f with vehicle (15% ethanol in sunflower seed oil). Mice remained in their cages for 6-7 weeks post-treatment to wash out potential tamoxifen effects before the time course collection.

Tissue collection was performed as described before.^14^ Briefly, mice were housed at 12:12 hour light/dark schedule. Before tissue collection, mice were housed in constant darkness for 30 hours. The mice were euthanized with cervical dislocation under dim red light, and heart tissue was collected every 4 hours for 48 hours. Tissues were flash-frozen for RNA and protein analysis. The recombination assay was assessed as described before.^14^

RNA isolation was performed as described before.^18^

Protein western blot was performed as described before.^19^

The Interdisciplinary Center for Biotechnology Research constructed an RNA sequencing library at the University of Florida. PolyA mRNA was isolated using the NEBNext poly(A) mRNA magnetic isolation module (E7490L), and the RNAseq library was prepared using the NEBNext Ultra II Directional RNA library Prep kit for Illumina (E7760L). We performed 150bp paired-end sequencing to a read depth for all samples at ∼ 40M reads.

RNA sequencing data and pathway analysis was performed as described before.^18^

Circadian rhythmicity analysis was performed as described previously using DiffCircadian.^18,20^

Differential expression analysis was performed by using DESeq2 in R studio.^21^ Significant DEGs were genes with an FDR < 0.05 and |log2FoldChange| ≥ 0.5.

## Results

### 1, RNAseq analysis reveals sex differences in the heart circadian transcriptome

To study the cardiac circadian transcriptome in male and female mice, total RNA was isolated from ventricular tissue collected every 4 hours for 48 hours under darkness (circadian times (CT) CT18 to CT62). The sequencing data was analyzed using DiffCircadian.^20^ In Table S1, we provide the number of rhythmically expressed genes (REGs) identified at different p-values or q-values. We identified more REGs in the female heart than the male heart at all p- and q-value cutoffs. For all downstream analyses, we defined REGs as those genes with a q < 0.05.

We first identified the size of the cardiac circadian transcriptome in the male and female mice. We identified 4644 REGs (33% of 13963 detectable transcripts) from the male hearts (Table S2) and 5678 REG (40.5% of 13980 detectable transcripts) from the female hearts (Table S3). To visualize the temporal expression pattern of those REGs, all REGs were sorted by their time of peak expression (peak time). As expected, the heatmap of z-score normalized expression over 24 hours from male (Figure 1A) and female REGs (Figure 1B) showed clear time-of-day rhythmic patterns.

**Figure 1.**
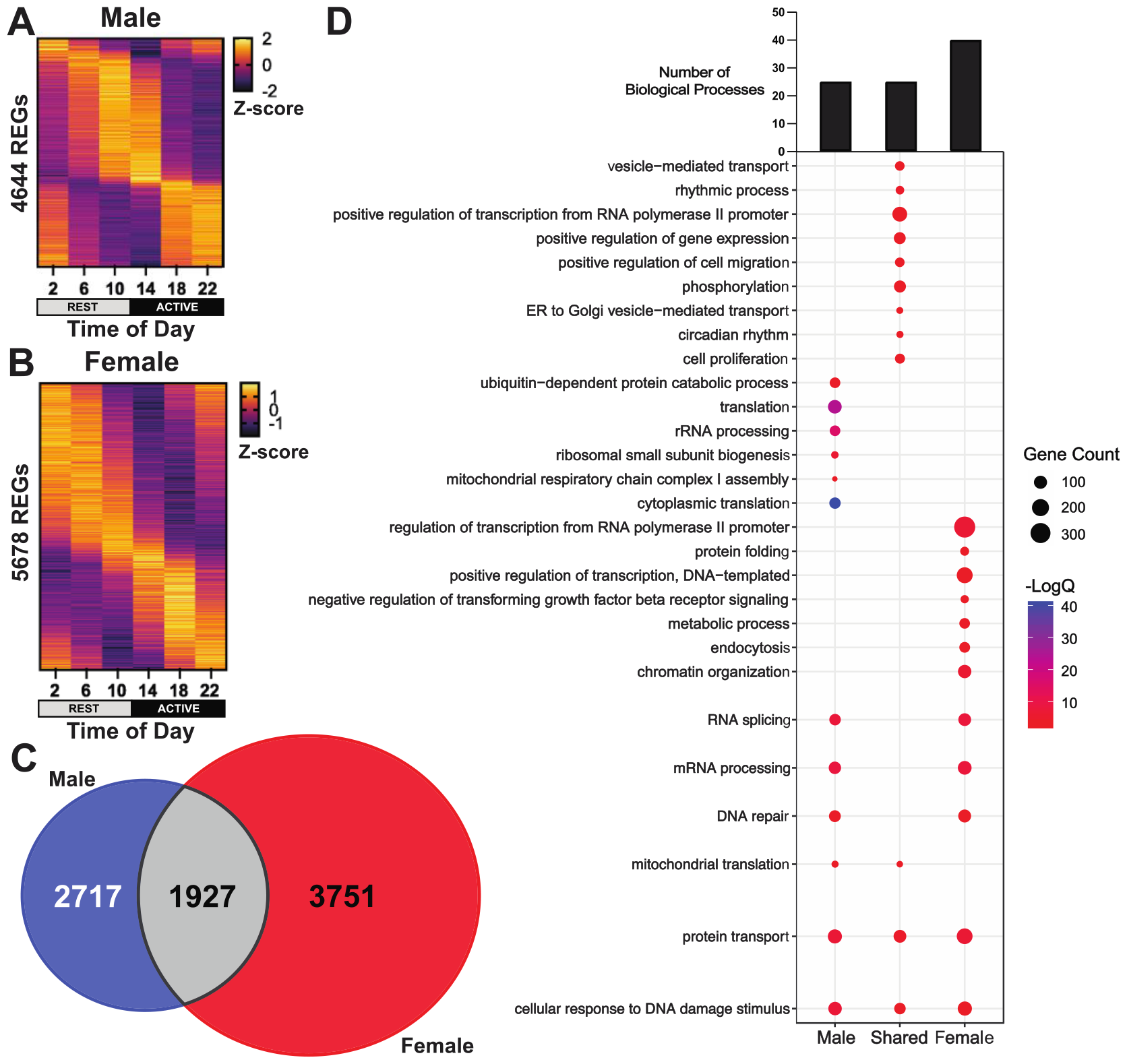
Comparison of circadian transcriptome between male and female mouse heart. Heatmap of z-scored REGs (the second day overlapped with the first day) with yellow as the highest expression level and brown as the lowest expression level for male mouse heart REGs (A) and female mouse heart REGs (B). (C) Venn diagram of REGs comparison between male (blue) and female (red) vehicle hearts with the gray for shared. (D). Enriched biological pathways. The total numbers of unique and shared biological pathways for each sex were shown on top of the bubble plot. The bubble plot shows some examples of the enriched pathways. Bubble size indicates the number of REGs enriched. The color indicates the logAdjQ value of enriched biological pathways, with red being the lowest and blue being the highest.

We then compared the male and female REGs and found ∼40% of the REGs (1927) were shared between males and females, with 3751 (66%) being female-specific and 2717 (58.5%) being male-specific (Table S4, Figure 1C). This level of overlap is much greater than the overlap between different tissues;^10^ however, it does indicate that ∼60% of REGs are sex-specific in the mouse heart.

We next asked whether the sex-specific REGs were enriched for similar or different functional pathways with gene ontology analysis performed using DAVID.^22,23^ We identified 25 enriched biological pathways from the male-specific REGs and 40 enriched biological pathways from the female-specific REGs (Figure 1D, Table S4) using an FDR < 0.05. Comparison of the enriched biological processes revealed eight common biological pathways (Figure 1D, Table S4), suggesting that circadian regulation of those biological processes does occur in both males and females, however it is through the rhythmicity of sex-specific genes. There were also biological pathways enriched by only male-specific REGs, such as translation, respiratory chain complex I assembly, ubiquitin-dependent protein catabolic process, and ribosome biogenesis (Figure 1D). In contrast, the female-specific REGs enriched chromatin organization, endocytosis, metabolic process, and protein folding (Table S4) (Figure 1D).

We then sorted all REGs by peak time and plotted the number of genes peaking across time of day (Figure 2A). We noted that the temporal distributions of male and female REGs were quite different. The majority of male REGs peak just before the transition of rest to active phase or active to rest phase. In females, the REGs also followed a bimodal distribution, but the peaks were broader and occurred in the middle of the rest and active phases. We next asked whether this pattern was isolated to just sex-specific REGs. As expected, the sex-specific REGs showed a bimodal distribution almost identical to the total REGs (Figure 2B). Interestingly, we also observed this same distribution pattern with the shared REGs (Figure 2C). These results demonstrate that the clock output is also sex-specific based on the temporal distribution of REGs throughout the day.

**Figure 2.**
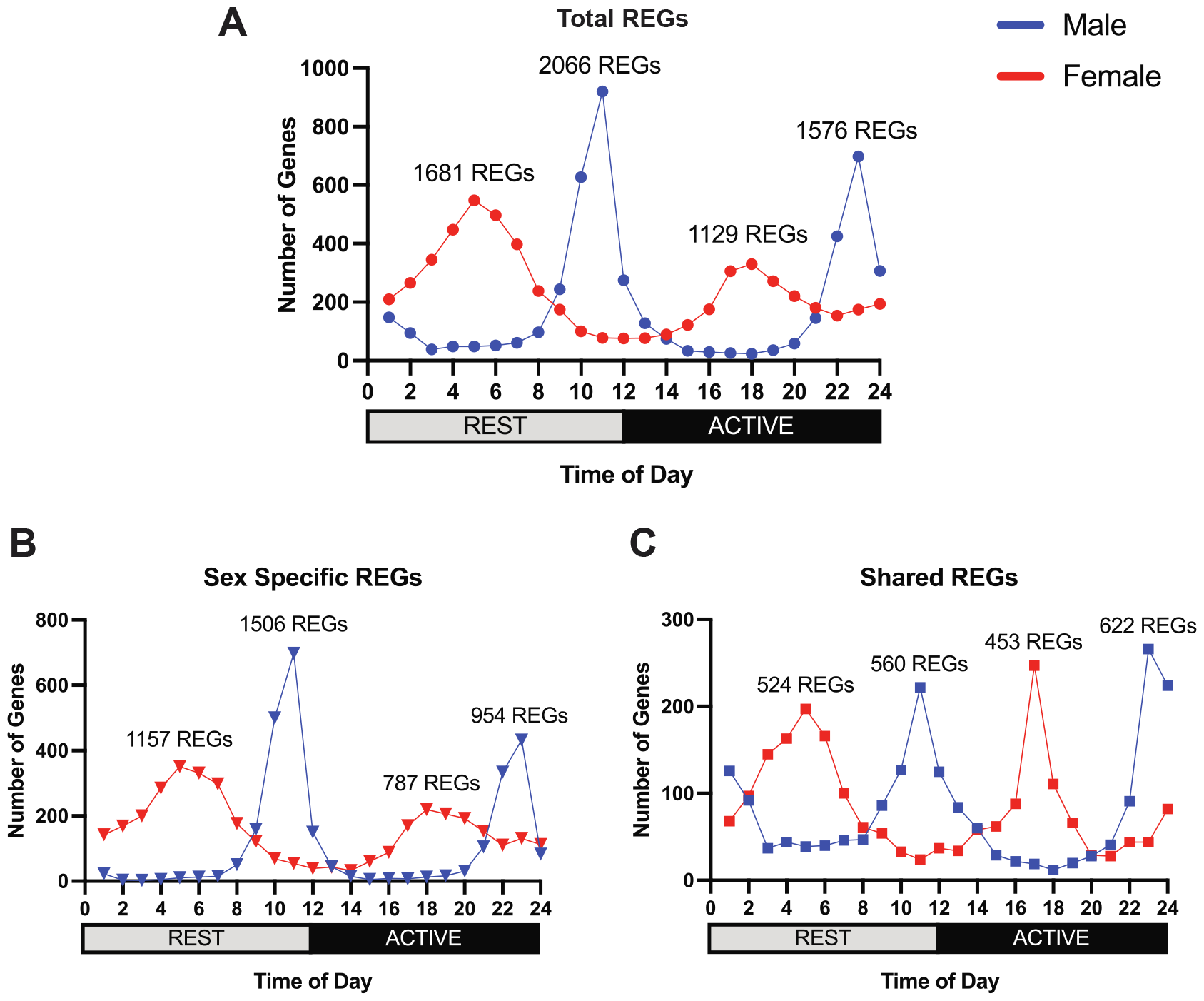
Phase distribution of male and female REGs over 24 hours. (A) phase distribution of all male (blue) and female (red) mouse heart REGs. (B) phase distribution of male-specific (blue) and female-specific (red) REGs. (C) phase distribution of male (blue) and female (red) shared REGs.

Next, we took the temporally distributed REGs and asked whether the pathways enriched by these REGs demonstrated any common temporal pattern. For this analysis, we binned all REGs into six 4-hour phases and subsequently performed gene ontology analysis to assign biological pathways to each phase (FDR < 0.05). As shown in Figure 3A, the biological pathways in the male heart have a similar phase distribution to that observed with male genes (Figure 2), largely clustering before activity onset and before activity offset. In females, the distribution of enriched biological pathways is largely consistent with the REGs but demonstrates slightly broader coverage over time-of-day (Figure 3B). This temporal analysis identified time-of-day-specific biological pathways shared between sexes, but interestingly, these shared pathways peaked in different phases of the day (Figure 3C). For example, protein transport, Golgi organization, and cellular response to DNA damage stimulus peaked in the middle of the rest phase in the female hearts vs. the end of the active phase in male hearts. Another cluster of shared biological pathways, including translation, mRNA processing, and mitochondrial pathways, peaked in the middle of the active phase for female hearts vs. the end of the rest phase in male hearts. These observations highlight two key points. First, there are some common biological pathways regulated by the clock mechanism in male and female hearts, but the timing of their expression appears offset in phase. Second, this analysis also identified sex-specific pathways such as autophagy in males and chromatin organization in females.

**Figure 3.**
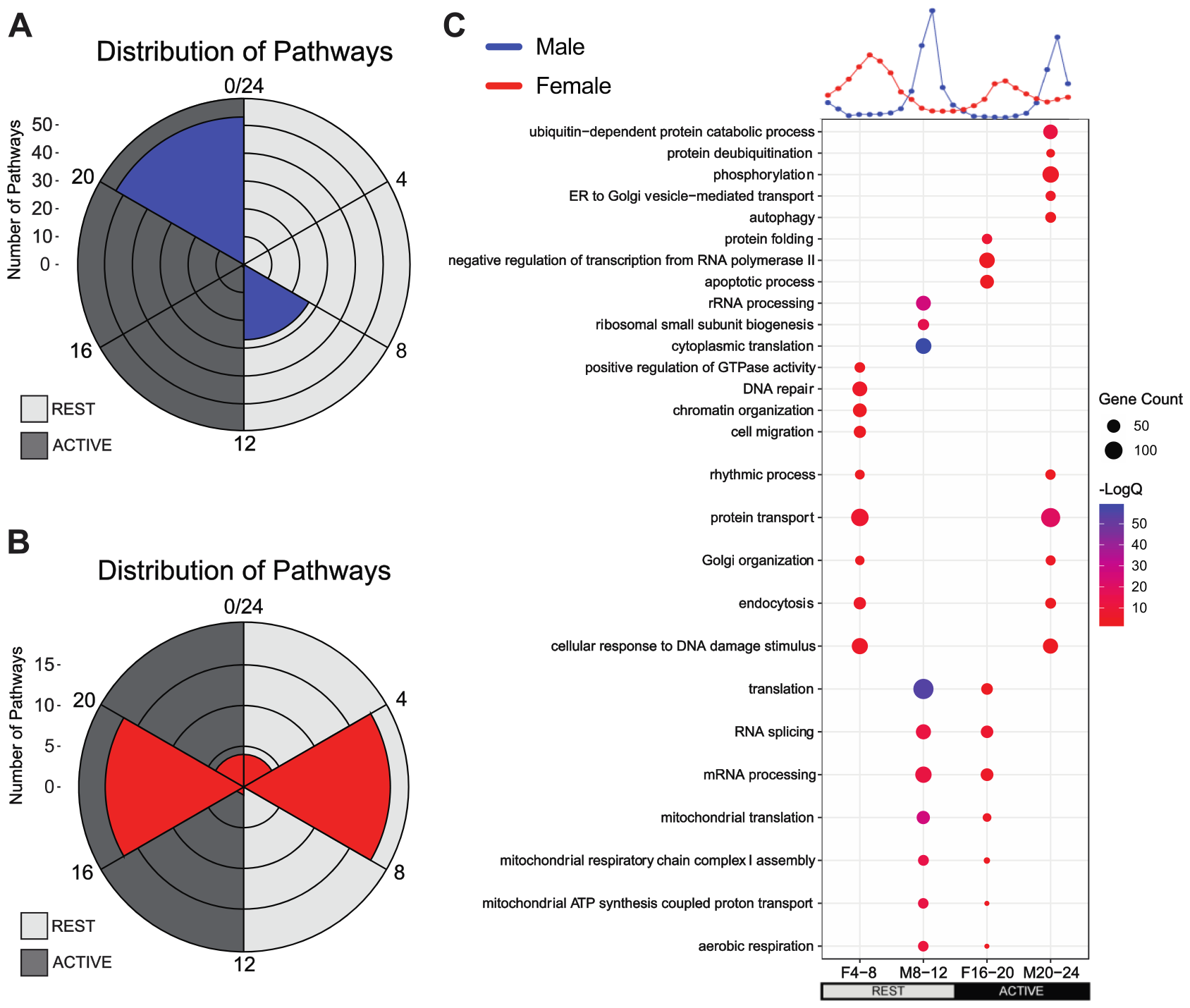
The temporal distribution of biological pathways. (A) phase distribution of enriched biological pathways of male hearts in 6 bins by circular plot. A gray background indicates a dark period, and a light background indicates a light period. (B) Phase distribution of female heart enriched biological pathways in 6 bins by circular blot. A gray background indicates a dark period, and a light background indicates a light period. (C) Enriched biological pathways. The phase distribution of REGs was shown on top of the bubble plot (red: female, blue: male). Each column represented the pathways enriched in each peak of the distribution (female peak from hours 4-8: F4-8, male peak from hours 8-12: M8-12, female peak from hours 16-20: F16-20, male peak from hours 20-24: M20-24). The bubble plot showed some examples of the enriched pathways. Bubble size indicates the number of REGs enriched. The color indicates the logAdjQ value of enriched biological pathways, with red being the lowest and blue being the highest.

We then queried whether the sex differences of the heart circadian transcriptome resulted from sex-specific expression of the core clock genes. Analysis revealed that the expression of all core clock genes, including *Bmal1, Clock, Per1, Per2, Cry1, Cry2, Nr1d1*, and *Nr1d2*, was almost identical between sexes (Figure S1 A). This observation indicates that sex-specific core clock expression cannot explain the observed differences in clock output.

### 2, *Bmal1* knockout in cardiomyocytes blunts the cardiac circadian transcriptome in male and female mice

To determine whether the molecular clock has a role in the sex differences of the cardiac circadian transcriptome in adult mice, we disrupted the cardiomyocyte molecular clock by generating an inducible cardiomyocyte-specific *Bmal1* KO mouse (iCS*Bmal1* KO). The recombination specificity was assessed by PCR as previously described, and diminished BMAL1 protein was assessed by western blots.^14^ We again performed a time course collection followed by RNA sequencing for this set of experiments as described above. Importantly, the iCS*Bmal1* KO mice do not show any signs of dilated cardiomyopathy at the age of collection.^14^ As shown in Figures S1B and S2, *Bmal1* mRNA and protein levels were blunted in male and female iCS*Bmal1* KO hearts. We suggest that the remaining *Bmal1* mRNA and protein expression are due to contributions from other cell types in heart tissue. As expected, we found that the expression of other core clock genes was significantly altered (Figure S1B). However, when we compared core clock gene expression between the male and female iCS*Bmal1* KO hearts, no significant differences were observed. These results confirmed that the cardiomyocyte core clock was significantly disrupted in our model, as described previously.^14^

Next, we analyzed the circadian transcriptome in the iCS*Bmal1* KO heart. As shown in Table S1, the number of REGs was significantly decreased in both male and female iCS*Bmal1* KO hearts compared to vehicle-treated hearts. We identified 1067 REGs (Table S5) in the male iCS*Bmal1* KO heart and 3641 REGs (Table S6) in the female iCS*Bmal1* KO heart, indicating that cardiomyocyte-specific *Bmal1*, and likely the clock mechanism, is required for the daily rhythmic expression of many genes in both the male and female heart tissues.

### 3, The circadian transcriptome of the male and female heart is altered in iCS*Bmal1* KO mice

In Figure 4A, we provide a heatmap of the rhythmic gene expression in the male iCS*Bmal1* KO heart. There is a large decrease in the number of REGs with loss of *Bmal1* with only 1067 REGs (∼77% decrease) identified in the male iCS*Bmal1* KO (Table S5). Of the 1067 REGs, we observed 583 uniquely rhythmic genes in the iCS*Bmal1* KO hearts, which were not rhythmic in the vehicle (Figure 4B and S3).

**Figure 4.**
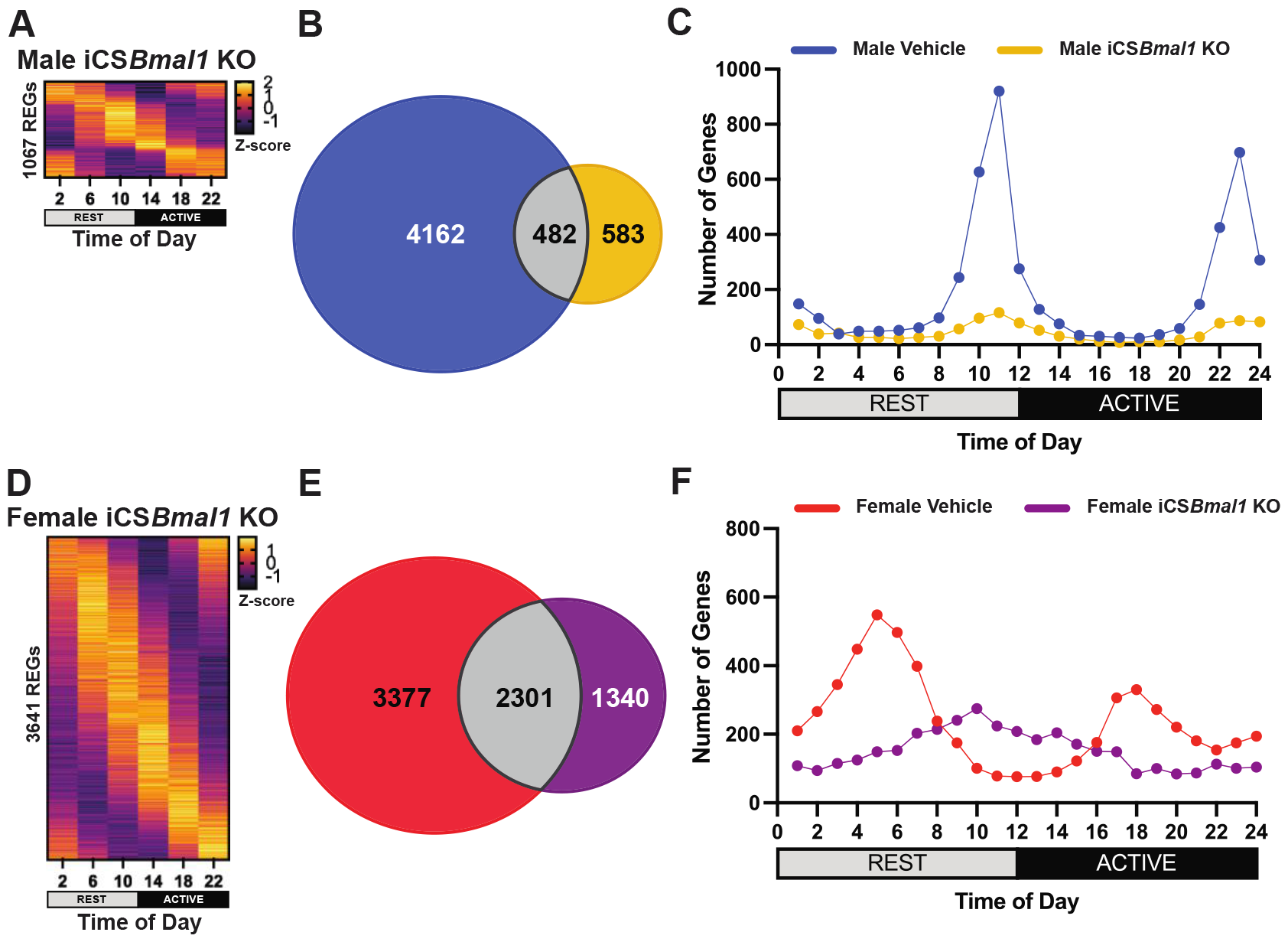
Significant changes to circadian transcriptome in male and female iCS*Bmal1* KO mouse hearts. (A) Heatmap of z-scored REGs in iCS*Bmal1* KO. Yellow is the highest expression level, and brown is the lowest expression level. (B) Venn diagram of REGs comparison between the vehicle (blue) and iCS*Bmal1* KO REGs (yellow) with shared in gray. (C) Comparison of the phase distribution of REGs over 24 hours between the vehicle (blue) and iCS*Bmal1* KO (yellow). (D) Heatmap of z-scored REGs in iCS*Bmal1* KO. Yellow is the highest expression level, and brown is the lowest expression level. (E) Venn diagram of REGs comparison between vehicle (red) and iCS*Bmal1* KO REGs (purple), with shared REGs in gray. (F) Comparison of the phase distribution of REGs over 24 hours between the vehicle (red) and iCS*Bmal1* KO (purple).

We then analyzed the temporal distribution of REGs in the male iCS*Bmal1* KO over 24 hours. Unlike the REGs in the male vehicle-treated hearts, the REGs in the iCS*Bmal1* KO hearts were more evenly distributed throughout the day (Figure 4C). The clusters of genes peaking before activity offset and onset in the male vehicle were significantly diminished in the iCS*Bmal1* KO hearts. This result indicates that cardiomyocyte-specific loss of *Bmal1* alters the overall temporal pattern of the REGs in male heart tissue with large-scale depletion of REGs that peak before activity onset and activity offset. Pathway analysis of the male iCS*Bmal1* KO REGs can be found in Figure S3 and Table S7.

Next, we examined the female cardiac circadian transcriptome in the iCS*Bmal1* KO mice. Surprisingly, we identified 3641 REGs (Table S6), only a ∼36% decrease compared to the vehicle-treated heart with the heatmap of expression showing a clearly rhythmic pattern over 24 hours (Figure 4D). We found that 63% (2301) of the REGs in the iCS*Bmal1* KO heart were shared with the vehicle-treated (Figure 4E). In contrast, the iCS*Bmal1* KO-specific REGs were not detected as rhythmic in the vehicle-treated heart but gained rhythmicity in the iCS*Bmal1* KO hearts (Figure 4E and S3A).

We then analyzed the temporal distribution of the REGs in the female iCS*Bmal1* KO heart over 24 hours. Like the changes in the male heart, we found that the two clusters of REGs observed in the middle of the rest phase and active phase in the female vehicle were no longer visible in the iCS*Bmal1* KO hearts (Figure 4F), leaving a broader peak of REGs that was more temporally aligned to the transition period from the rest phase to the active phase. Pathway analysis of the female iCS*Bmal1* KO REGs can be found in Figure S4 and Table S8.

### 4, Sex differences in the heart circadian transcriptome are diminished in iCS*Bmal1* KO mice

Lastly, we asked whether loss of *Bmal1* resulted in changes to the sex-specific gene expression observed in the vehicle-treated hearts. As seen in Figure 5A, we found 752 REGs out of a total of 1067 male iCS*Bmal1* KO REGs were shared with the female iCS*Bmal1* KO REGs, representing a 70% overlap, much larger than that of the vehicle-treated hearts. Gene ontology analysis identified 14 enriched biological pathways from the shared 752 REGs and 40 enriched biological pathways from the female-specific REGs, with no pathways enriched by the male-specific REGs (Table S9, Figure 5B). Half of the biological processes enriched by shared REGs were also found to be enriched by female-specific REGs, including cellular response to DNA damage stimulus, rhythmic process, DNA repair, positive regulation of transcription from polymerase II, mRNA processing, and protein phosphorylation, among others (Figure 5B). These observations indicate that loss of cardiomyocyte-specific *Bmal1* results in a greater relative overlap in the REGs between males and females. In addition, the large number of REGs in the female iCS*Bmal1* KO heart suggests a greater resilience and/or contribution of other non-cardiomyocyte cells to the cardiac tissue circadian transcriptome in females.

**Figure 5.**
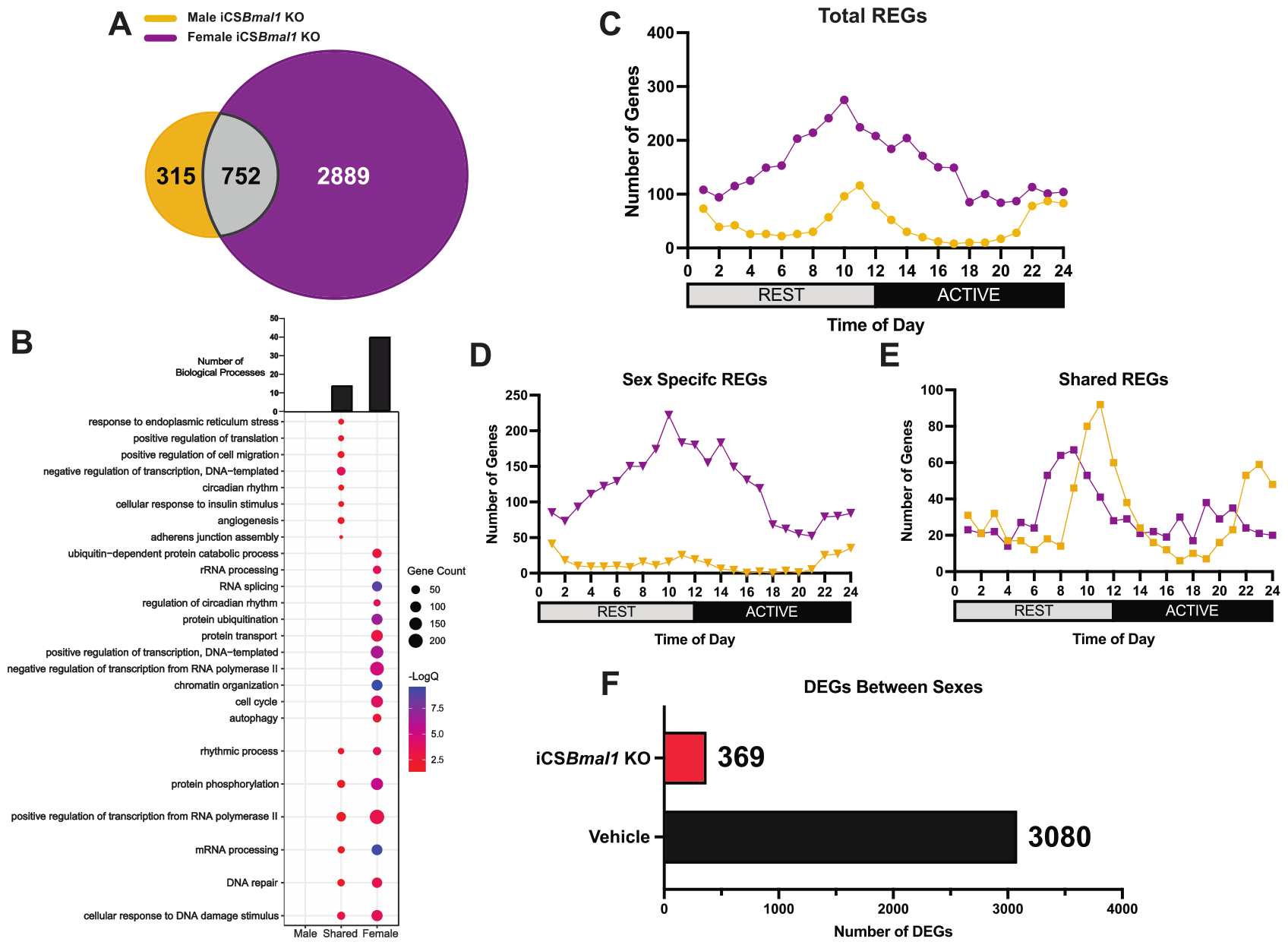
Comparison of male and female iCS*Bmal1* KO circadian transcriptomes. (A) Venn diagram of REGs comparison between male (yellow) and female (purple) iCS*Bmal1* KO hearts, with shared REGs in gray. (B) Enriched biological pathways. The total number of unique and shared biological pathways between male and female iCSBmal1 KOs was shown on top of the bubble plot. The bubble plot showed some examples of the enriched pathways. Bubble size indicates the number of REGs enriched. The color indicates the logAdjQ value of enriched biological pathways, with red being the lowest and blue being the highest. (C) Comparison of the phase distribution of REGs of male (yellow) and female (purple) iCS*Bmal1* KO. (D) Comparison of the phase distribution of sex-specific REGs of male (yellow) and female (purple) iCS*Bmal1* KO. (E) Comparison of the phase distribution of shared REGs between male (yellow) and female (purple) iCS*Bmal1* KO. (F) Bar graph of differential gene expression between male and female heart transcriptome before and post-iCSBmal1 knockout. Black: vehicle-treated DEGs; red: iCS*Bmal1* KO DEGs.

Next, we compared the phase distribution of the male and female iCS*Bmal1* KO REGs. As shown in Figure 5C, the distribution of REGs in the male and female iCS*Bmal1* KO hearts was more similar than what was seen in the vehicle-treated hearts (Figure 2A), with the males exhibiting smaller peaks at the transition times while the female iCS*Bmal1* KO REGs were more broadly distributed across the day with a generalized peak over the transition from rest to active phase. As done previously, we looked at the distribution of the sex-specific and shared REGs. With the sex-specific REGs (Figure 5D), the sex difference was driven primarily by the number of REGs rather than the time-of-day pattern. With the shared REGs (Figure 5E), the female REGs exhibited a peak in the number of REGs at a slightly advanced phase compared to the male, but overall, the pattern was much more similar. When compared to those obtained with the vehicle-treated REGs, these patterns demonstrate a significant loss of sex differences in the cardiac circadian transcriptome after cardiomyocyte-specific deletion of *Bmal1*.

Our initial hypothesis was that the diminished sex differences with iCS*Bmal1* KO would be confined to the REGs and would not impact sex-specific expression of non-rhythmic genes. To address this concept we performed differential gene expression analysis with DESeq2^21^ using our dataset. Gene expression at each time point was treated as a replicate within a given genotype to assess differential expression independent of the time of day. In the vehicle-treated mice, we identified 3080 DEGs between male and female hearts (Table S10). In contrast, in the iCS*Bmal1* KO mice, we observed only 369 DEGs between sexes, illustrating an over 8-fold decrease compared to the vehicle (Figure 5F). This dramtic drop in sex-specific gene expression in the hearts was striking and argues that *Bmal1* in cardiomyocytes alone is largely responsible for sex-specific gene expression in the mouse heart. These data corroborate the observations made above; the cardiac transcriptome becomes less different between sexes following iCS*Bmal1* KO, indicating that the cardiomyocyte circadian clock is largely influential in the sex-specific gene expression in the heart.

## Discussion

In this study, we observed sex differences in the cardiac circadian transcriptome in mice. This was apparent in the quantity, identity, and temporal distribution of REGs; the biological processes enriched by those REGs; and the magnitude of differential expression between sexes. Of particular interest is the timing of peak REG expression between sexes as seen in Figure 2; even REGs shared by both males and females display a sex-specific temporal distribution throughout the day. While differential rhythmicity in gene expression has been observed between males and females in the past,^17^ we provide the first observation of a ∼8-hour phase difference in peak REG expression between sexes.

Following cardiomyocyte-specific disruption of the molecular clock, the number of REGs identified in both males and females decreased. However, circadian gene expression appears to be more resilient to clock disruption in females, similar to what has been observed in other tissues.^15^ Interestingly, the sex-specific temporal patterns in the distribution of cardiac REGs became visually less different following *Bmal1* knockout. Additionally, iCS*Bmal1* KO led to an over 8-fold decrease in the magnitude of differential gene expression between sexes. These data indicate that the cardiomyocte is the primary source of sex-specific gene expression in cardiac tissue and that the circadian clock is essential to this sex difference. Our results also suggest that the other cell types within the heart exhibit significantly less sex differences in gene expression.

While the cardiac circadian transcriptome exhibits sex differences, there is still very little understood about the mechanism of this difference. These results cannot be explained by differences in the expression of the core clock, as we did not find significant differences in core clock gene expression between sexes. This has also been observed recently in mouse liver,^16^ as well as in studies using male and female human samples from multiple tissues.^17^ It has been shown that commonly expressed transcription factors are not differentially expressed between males and females, but rather display sex-specific activity.^6,7^ This work has put forward that the transcriptional differences between different cell types and sexes are not just driven by a few specific transcription factors but by an organized transcription factor network.^24^ Therefore, *Bmal1* and the molecular clock components may have sex-specific binding partners and interact with larger regulatory networks that influence DNA binding stability, orientation, and spacing, resulting in differential gene expression between sexes.^25–27^ Thus, the sex differences we observed could be mediated by interactions between the clock and other transcription factor networks.^24,28,29^

Differential rhythmicity of other clock interacting factors may influence circadian gene expression between sexes. One example is SIAH2 ubiquitin ligase, a female-specific regulator of circadian rhythm. Previous work has shown that deficiency of *Siah2* in females, but not in males, resulted in alteration of core clock gene expression and remodeling of the liver circadian transcriptome by flipping the expression of rhythmic genes from the dark phase to the light phase.^30^ In our analysis, *Siah2* was identified as rhythmically expressed in females (q = 0.000869) but not rhythmic in males (q = 0.082). The differential rhythmicity of *Siah2*, in addition to its sex-specific role in rhythm maintenance, may be indicative of differences in the larger regulatory networks that contribute to sex-specific clock output.

Some other possible factors that could impact sex differences include sex hormone receptors and sex chromosomes.^31^ It has been shown that the differential presence of androgens between sexes influences the magnitude of gene expression in other tissues.^32^ The possible cooperative interaction between clock components, sex hormones, and their receptors could mediate the differential targeting and activation of circadian gene expression. However, sex hormone receptor motifs are not enriched by DEGs between sexes.^6^ Similarly, we did not find sex hormone receptor motif enrichment in the regulatory regions of the REGs in either the male or female cardiac circadian transcriptome (data not shown). However, we did identify REGs located on the X chromosome in both the male and female circadian transcriptome but did not find any REGs on the Y chromosome. Similar to what we observed in the entire circadian transcriptome, the phase distribution of REGs from the X chromosome exhibited clear sex differences (data not shown). Further study is required to delineate the role of the X chromosome on the sex differences of the circadian transcriptome.

In conclusion, differential gene expression between sexes has been shown to contribute to sexually dimorphic phenotypes and diseases.^6^ Here, we identified sex differences in the circadian transcriptome in mouse hearts and showed that cardiomyocyte *Bmal1*, and likely the core clock mechanism, is indispensable for these sex differences. These findings provide new insights into mechanisms directing sex-specific gene expression programs. In addition, they provide important reference data to support our understanding of sex differences in health and disease.

